# Whole-genome benchmarking reveals context-specific error rates in the Ultima UG100 and Illumina NovaSeqX Platforms

**DOI:** 10.64898/2026.02.27.708547

**Authors:** Oona Shigeno Risse-Adams, Paul Collier, Theodore M Nelson, Jonathan Foox, Christopher E Mason

## Abstract

Whole genome sequencing (WGS) is becoming more common in research and clinical applications, but there is a paucity of comparative data between high-throughput WGS platforms. We benchmarked the Ultima Genomics UG100 system against Illumina NovaSeqX using the Genome in a Bottle (GIAB) WGS reference set sample HG002. Across NIST v4.2.1 benchmark regions, UG100 exhibited higher total variant-calling errors (27x), primarily driven by indel false negatives. Restricting to Ultima Genomics’ high confidence regions (UG HCR v3.1, 90.3% of GRCh38) reduced the error burden by 89.6%, indicating most errors lie outside these regions. Strong degradation of variant calling performances occurred in homopolymer tracts > 10 bp, and base calling error rates increased sharply after position 200 bp within each read, whereas Illumina’s error rates were higher earlier in the read. Coverage dropouts in UG100 data were pronounced in GC-rich regions. Of note, 2.24% of ClinVar Pathogenic and Likely Pathogenic variants and 22.6% of a common catalogue of polymorphic STRs were found to be excluded from UG HCR v3.1, indicating potential impact to clinical applications. These results highlight context-specific genotyping errors of the UG100 and NovaSeqX platforms, and underscore the importance of whole genome benchmarking for adjudicating accuracy of NGS platforms beyond the current HG002 reference set.

## Introduction

Benchmarking new short-read platforms requires transparent evaluation against community standards, and is also critical for improvements in Next-Generation Sequencing (NGS) and clinical applications. We compared Ultima Genomics UG100 with Illumina NovaSeqX using the GIAB (NIST v4.2.1) framework ^1,2^ on HG002 (GRCh38) ^3^. For a comprehensive assessment, we checked accuracy in the full NIST benchmark and within Ultima Genomics defined UG High Confidence regions (UG HCR v3.1). We then evaluated sequence-context effects (homopolymer length), read-cycle error behavior, replicate reproducibility, and potential clinical impact via GC/STR analyses. Our results show a higher UG100 error burden compared to the NovaSeqX system, with variants concentrated in specific contexts, and largely absent when restricting to UG HCR.

## Methods

### Study Design

We evaluated HG002 (NA24385; GRCh38) on NovaSeqX 25B v1.3 and UG100. Data were processed with standardized, platform-appropriate pipelines (NovaSeqX: DRAGEN Germline v4.4 ^5^; UG100: Ultima AWS Ready-to-Run DeepVariant v1.0 ^4^). Datasets were matched to ∼35x target raw coverage (including duplicate reads), and variant calling accuracy was assessed against GIAB NIST v4.2.1 truth VCF/BED ^1^. We report (i) whole-benchmark accuracy; (ii) accuracy restricted to UG HCR v3.1; and (iii) context-stratified accuracy (by homopolymer length).

### Datasets

Seven independent Ultima Genomics UG100 runs ^6^ were generated at the New York Genome Center (NYGC) using NEB PCR-free library preparation. Two runs employed the ppmSeq chemistry, while five runs used the standard WGS chemistry. No Solaris chemistry was used. Samples included the GIAB HG002 reference genome (three technical replicates in each run with standard WGS chemistry; five technical replicates in each run with ppmSeq chemistry). All libraries were PCR-free, and data were delivered as aligned CRAM files.

For Illumina NovaSeq, replicates of GIAB HG002 reference genome were prepared using TruSeq DNA PCR-Free library prep kit as part of a 64-plex pool. The libraries were sequenced on 3 runs with NovaSeqX 25B reagents using 2x151bp read length and the NovaSeqX Sequencing System Software Suite v1.3.0. One HG002 library from each run is included in this study.

### Data hosting and access

All sequencing data generated in this study have been deposited in the NCBI Sequence Read Archive (SRA). See Data Availability section for accession numbers.

### Reference and Annotation Resources

Analyses used the GRCh38 reference genome ^3^, the GIAB NIST v4.2.1 truth VCF and BED ^1^, Ultima High Confidence Regions (v3.1), ClinVar variants (release 2024-08-05) ^7^, a genome-wide Tandem Repeats catalog ^8^, and in-house homopolymer annotations generated from GRCh38.

### Read-length Estimation

Read-length statistics were computed for Ultima Genomics UG100 CRAMS using samtools v1.15.1 ^9^. Full arguments and scripts are provided in supplementary note 1. For UG100 runs with ppmSeq chemistry, we additionally computed read lengths separately for duplex reads. To achieve this, we extracted duplex reads from the original CRAM files by selecting all reads where both st and et tag were set to “MIXED”. Distributions were summarized across all replicates.

### Base-calling Error Estimation

To quantify base-calling error rates, we analyzed aligned reads (MAPQ > 30) within NIST v4.2.1 high confidence regions (HCR). Known variants from the truth VCF were masked to prevent counting true polymorphisms as errors. For each base, we compared the sequenced nucleotide to the reference and tallied mismatches, insertions, and deletions, stratifying by predicted Q-score, read position, and read pair (R1/R2). We reported error rates by read position separately for substitutions and for indels (combined insertions and deletions).

For UG100 runs with ppmSeq chemistry, we additionally computed base-calling error rates separately for duplex reads. To achieve this, we extracted duplex reads from ppmSeq datasets as described in the previous paragraph. Finally, for all replicates we reported substitution error rates restricted to the bases with the highest predicted Q-score bin (Q35 for UG100 and Q40 for NovaSeqX). For improved readability, we reported error rates on a phred-scale as empirical Q-scores. This metric highlights the level of error that is empirically detected in the bases that each platform reports as top quality. Average percentage of bases reported in the Q40 bin was 94.22% for NovaSeqX 25B runs. Average percentage of bases reported in the Q35 bin was 84.33% for UG100 runs with standard chemistry and 87.75% for UG100 runs with ppmSeq chemistry.

### Downsampling, Alignment and Variant Calling

#### NovaSeqX

FASTQs were aligned and variants called using the Illumina DRAGEN Germline Pipeline ^5^ v4.4 with the Map/Align + Small Variant Caller module (with personalization activated). Downsampling to 35x target raw coverage (including duplicate reads) was performed using fractional subsampling.

#### Ultima Genomics UG100

In UG100 runs, sequencing from individual libraries did not provide sufficient depth to reach the standard 30x-40x coverage typically required for whole-genome sequencing assays. Raw aligned CRAM files from two libraries were therefore downsampled to ∼18x coverage using samtools v1.21 ^9^ and merged to reach 36x raw coverage (including duplicate reads). The same downsampling strategy was employed for datasets generated with standard chemistry and with ppmSeq chemistry. For UG100 datasets with ppmSeq chemistry, we additionally selected duplex reads only (as described above) and downsampled to 36x raw coverage with the same method. Variant calling for all datasets employed the Ultima AWS Ready-to-Run DeepVariant workflow (v1.0) ^4^.

#### Variant-calling benchmarking

Benchmarking was performed with hap.py v0.3.15 ^10^ by comparing each callset to the NIST truth VCF. Two stratification types were evaluated:

1. NIST v4.2.1 benchmark regions (full-genome benchmark)
2. NIST v4.2.1 benchmark regions restricted to Ultima High Confidence Regions v3.1 and their complement (UG Low Confidence Regions)

Complement beds were generated using bedtools v2.32.0 ^11^. Per-region Type 2 accuracy FP, FN, and F1 metrics were extracted from the extended CSV output for visualization.

#### Homopolymer Resolution Identification and Stratification

Homopolymer tracts ≥ 2 bp were identified across GRCh38 as contiguous runs of identical bases in the reference fasta. Output annotations in bed file format were stratified by homopolymer length (2,3,4…20 bp and > 20 bp).

#### Variant-calling accuracy by homopolymer length

For each homopolymer stratum, variant calls were benchmarked against the NIST v4.2.1 truthset using hap.py v0.3.15 ^10,12^. Type 2 accuracy true positives (TP), false positives (FP), and false negatives (FN) were summarized by homopolymer length. Indel precision and recall was plotted against homopolymer size to assess resolution limits (Fig. 3).

#### Reproducibility across replicates

For HG002 replicates variant overlaps were computed with bedtools multiinter ^11^ to estimate variant calling reproducibility. For NovaSeqX, two additional runs on 10B flowcells were used in this assessment to match the number of runs executed on UG100. UG100 datasets with ppmSeq chemistry were excluded from this assessment, as only two sequencing runs were available. Variants were considered to be reproducible if present at the same location in VCFs from multiple sequencing runs and attributed a PASS filter. Analyses were repeated genome-wide and restricted to NIST v4.2.1 HCR to compare intra-platform reproducibility of SNP and indel calls.

### Clinical impact assessment

#### ClinVar pathogenic variant overlap

Pathogenic (P) and likely pathogenic (LP) variants were extracted from the ClinVar 2024-08-05 release ^7^ using bcftools v1.22 ^9^. Overlaps between ClinVar P + LP sites and UG HCR v3.1 were quantified with bedtools intersect, and the proportion of clinically annotated bases excluded from the UG HCR was reported.

#### Tandem-repeat catalog overlap

UG HCR v3.1 was intersected with a genome-wide Tandem Repeats catalog ^8^ to estimate the fraction of polymorphic repeat loci outside UG HCR v3.1 regions.

## Results

Within the GIAB/NIST v4.2.1 benchmark regions, Ultima Genomics UG100 produced 44,593 SNP and 108,076 indel combined false positive and false negative calls (median values across replicates). In contrast, NovaSeqX exhibited roughly one-twenty-seventh as many total errors at matched coverage. The excess of UG100 was dominated by indel false negatives, consistent across technical replicates.

UG100 ppmSeq datasets yielded higher variant calling errors compared to UG100 datasets with standard chemistry for SNP false negatives (1.45 median fold increase), indel false positives (1.72 median fold increase), indel false negatives (1.38 median fold increase). Only SNP false positive errors were reduced (0.9 median fold decrease). Similar trends were observed in datasets restricted to duplex reads only.

The UG HCR region spans 90.30% of GRCh38 by interval length (computed from bed), and we define UG Low Confidence Regions (UG LCR) as (i) the whole-genome complement for descriptive summaries and (ii) the NIST-restricted complement (NIST v4.2.1 minus UG HCR) for benchmarking within the GIAB confidence regions. This leaves 99.07% of NIST v4.2.1 high-confidence regions and 93.2% of NIST v4.2.1 true variant calls within UG HCR. Of those remaining variants, found that 2.24% of ClinVar pathogenic and likely pathogenic variant sites are not included in UG HCR v3.1.

In addition, to evaluate the overlap of Ultima Genomics high confidence regions (UG HCR v3.1) with polymorphic tandem repeats loci, we utilized a publicly available genome-wide database (the Tandem Repeats catalog), a resource that contains 49 million loci of STRs and annotated tandem loci. We found that 22.58% of bases in the catalog (14,832,550 bps out of 65,678,112 bps) are not included in UG HCR v3.1.

## Discussion

Ultima Genomics UG100 currently has substantially higher whole-genome variant-calling error rates than Illumina NovaSeqX 25B flowcells, when both are run at matched 35x raw coverage on HG002 and evaluated against the GIAB NIST v4.2.1 truth set ^1^. Across NIST benchmark regions, UG100 has roughly 27-fold more total errors (false positives plus false negatives) than NovaSeqX, with the excess driven primarily by indel false negatives. This difference persists despite using each vendor’s recommended pipeline (DeepVariant ^4^ for UG100 and DRAGEN v4.4 ^5^ for NovaSeqX), and is more consistent with platform-specific read properties and error modes than with an obvious misconfiguration.

Ultima’s High-Confidence Regions (UG HCR v3.1) span about 90% of GRCh38 and include over 99% of NIST v4.2.1 benchmark intervals, yet restricting evaluation to these regions removes nearly 90% of UG100 variant calling errors. In other words, most of UG100’s error burden is concentrated in designated “Low Confidence” regions. From a user standpoint, summary metrics restricted to UG HCR understate true genome-wide variant calling error rates, but this information is critical for ensuring a full-genome evaluation of the accuracy ofhigh-throughput NGS platforms.

Our analyses additionally make clear that UG100’s errors are not uniformly distributed, but are context dependent. Indel performance on UG100 declines sharply as homopolymer length increases, with particularly poor recall in homopolymers of 20 bp or longer, whereas NovaSeqX maintains nearly constant performance across the same bins. Roughly two-thirds of UG100 variant-calling errors occur in homopolymers of at least 10 bp. Error-by-cycle profiles and read-length distributions help explain this behavior: UG100 substitution error rates are relatively low through the first ∼200 cycles but increase towards the read tail, and the median read length of roughly ∼300 bp places many bases in this high-error region, which indicates that read-trimming could be a method of improving these errors raters. Of note, indel error rates are elevated throughout the read length, and so trimming may not be as effective on those contexts. The ppmSeq chemistry, especially when restricting to duplex reads, shortens effective read length and reduces overall error levels, but the late-cycle error increase remains.

Beyond homopolymers and read tails, UG100 also shows more pronounced GC bias and coverage dropout than NovaSeqX, especially above ∼60% GC. NovaSeqX coverage is comparatively flat across GC bins, whereas UG100 coverage declines in GC-rich sequence and drops markedly at the highest GC values. When considering clinically relevant loci, a non-trivial fraction of ClinVar pathogenic and likely pathogenic variants, as well as a substantial proportion of bases in a genome-wide tandem-repeat catalog, fall outside UG HCR, including high-GC short tandem repeat loci such as the FMR1 CGG repeat. Reproducibility analyses show a similar pattern: while SNP calls are highly reproducible on both platforms in NIST high-confidence regions, indel calls on UG100 are consistently less reproducible than on NovaSeqX, particularly when considering whole-genome calls. These features could complicate cohort wide analyses and longitudinal studies, where consistent indel calls in homopolymer rich and GC-rich regions are often critical.

This study has limitations. We assessed only a single sample (HG002), one Illumina configuration (NovaSeqX 25B with DRAGEN v4.4 germline ^5^), and one Ultima configuration (UG100 with the AWS Ready-to-Run DeepVariant v1.0 workflow). We did not evaluate Solaris chemistry, alternative UG-specific callers, or customized variant-calling pipelines. Our benchmarking is restricted to NIST v4.2.1 truth-set regions and their overlap with UG HCR, leaving regions outside NIST largely uncharacterized for both platforms, and it is notable that Illumina software has extensively trained on HG002. Future chemistry updates, base-calling models, or variant callers may narrow the gap we observe.

Despite these caveats, under a common GIAB/NIST benchmarking framework ^1,13^ at matched coverage, UG100 currently exhibits higher genome-wide variant-calling error rates than NovaSeqX, with errors concentrated in long homopolymers and late read cycles, and with coverage and confidence deficits in high-GC regions, many of which fall outside the vendor’s high-confidence genome regions. Conversely, Illumina exhibits a higher error rate earlier in the read. For applications that require reliable whole-genome coverage, such as clinical sequencing, rare-variant discovery, and interpretation in regulatory, repetitive, or GC-rich regions, these context-specific limitations need to be explicitly considered. More broadly, our results argue for community-standardized, sequence-context-aware benchmarking of sequencing platforms ^3,12^. The framework used here provides one template for such evaluations and highlights the risk of over-interpreting global precision/recall statistics or metrics restricted to selected confidence regions when judging clinical or whole-genome readiness.

## Figures

**Figure 1.**
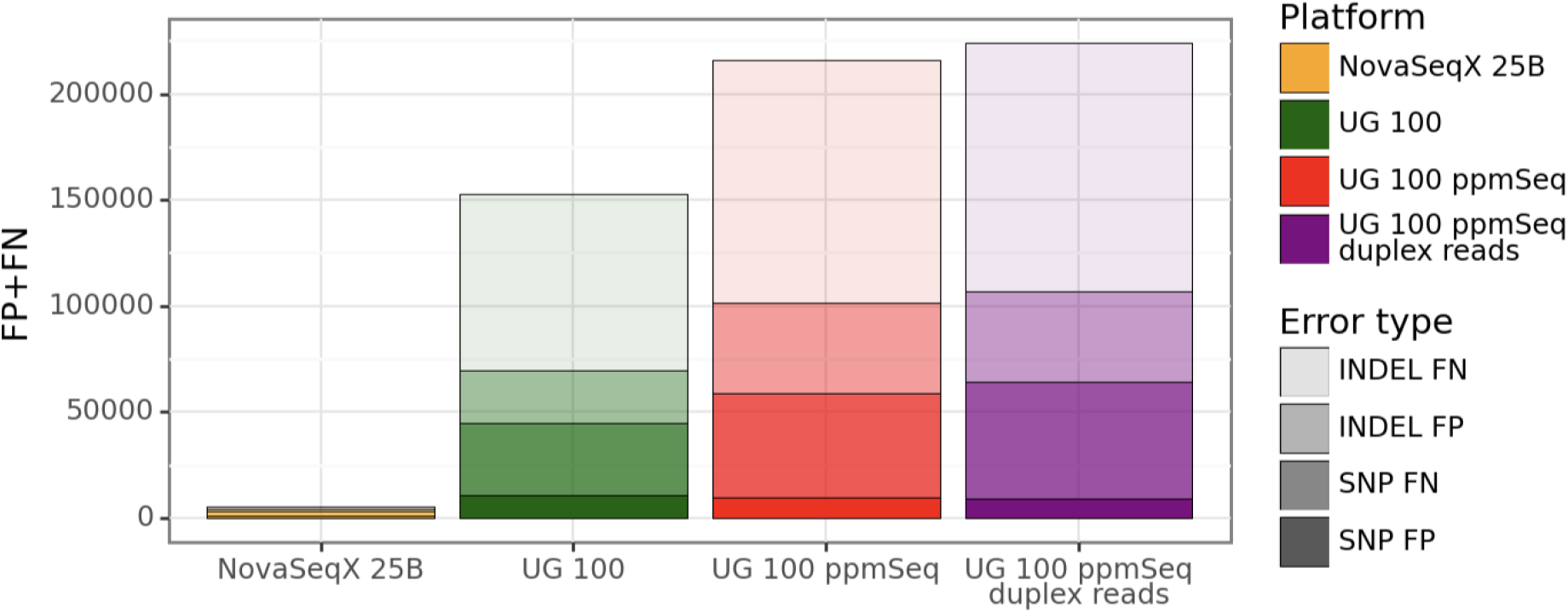
Whole-genome variant calling accuracy gap between UG100 and NovaSeqX. Indel and SNP False Positive and False Negative errors (FP+FN) within NIST v4.2.1 are shown for NovaSeqX, UG100, UG100 ppmSeq, and UG100 ppmSeq duplex reads (median values across replicates), stratified by variant and error type. Against NIST v.4.2.1 regions, UG100 showed a median 27-fold higher combined total variant-calling error burden relative to NovaSeqX. Variant calling errors in UG100 ppmSeq datasets (all reads or duplex reads only) were higher compared to UG100 across all variant and error types, with the exception of SNP False Positives, which decreased.

**Figure 2.**
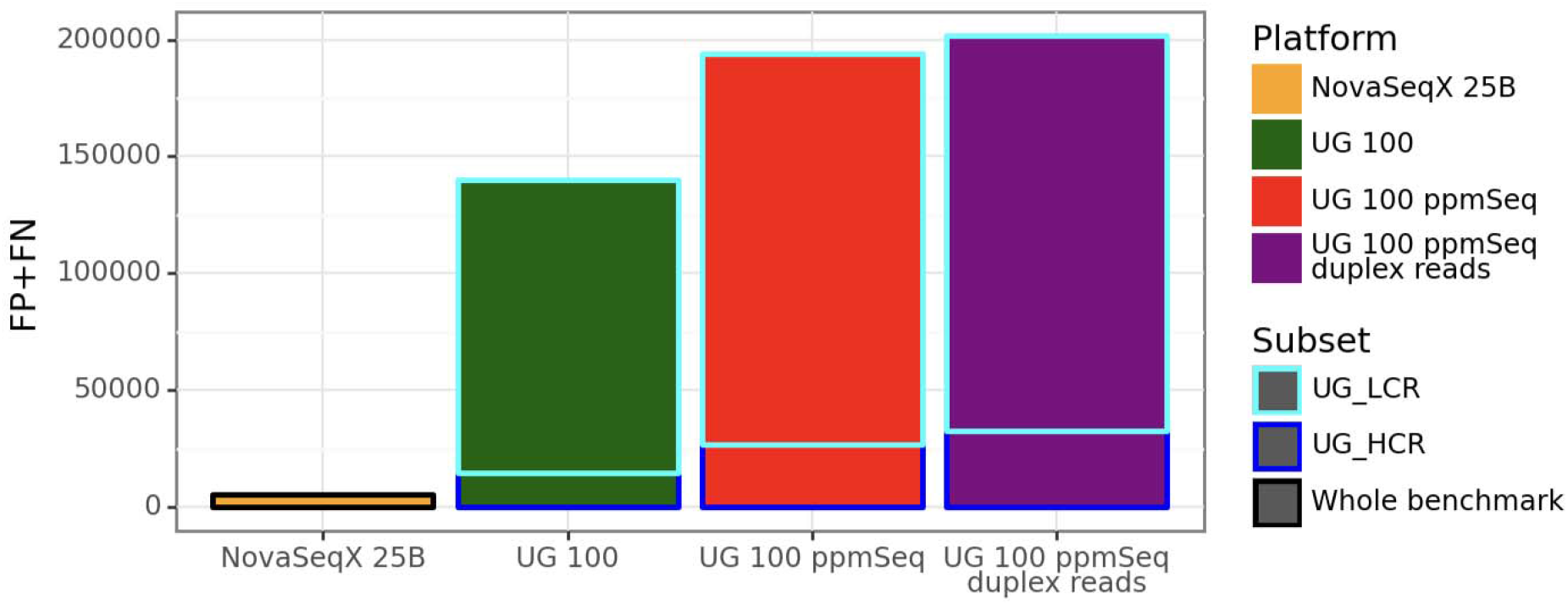
Most Ultima Genomics UG100 errors occur outside the vendor’s High-Confidence Regions. Indel and SNP False Positive and False Negative errors (FP+FN) within NIST v4.2.1 are shown for NovaSeqX 25B, UG100, UG100 ppmSeq, and UG100 ppmSeq duplex reads (median values across replicates). Restricting to UG HCR v3.1 (90.3% of GRCh38; 99.07% of NIST v4.2.1 high-confidence regions; 93.2% of NIST v4.2.1 truth variant calls) removes 89.6% of UG100 FP+FN, indicating the majority of UG100 errors lie in vendor-defined low-confidence regions. The same trend is observed in UG100 ppmSeq datasets, with 86.2% of variant calling errors in UG LCR v3.1 in ppmSeq datasets and 83.9% of variant calling errors in UG LCR v3.1 in ppmSeq datasets restricted to duplex reads only.

**Figure 3.**
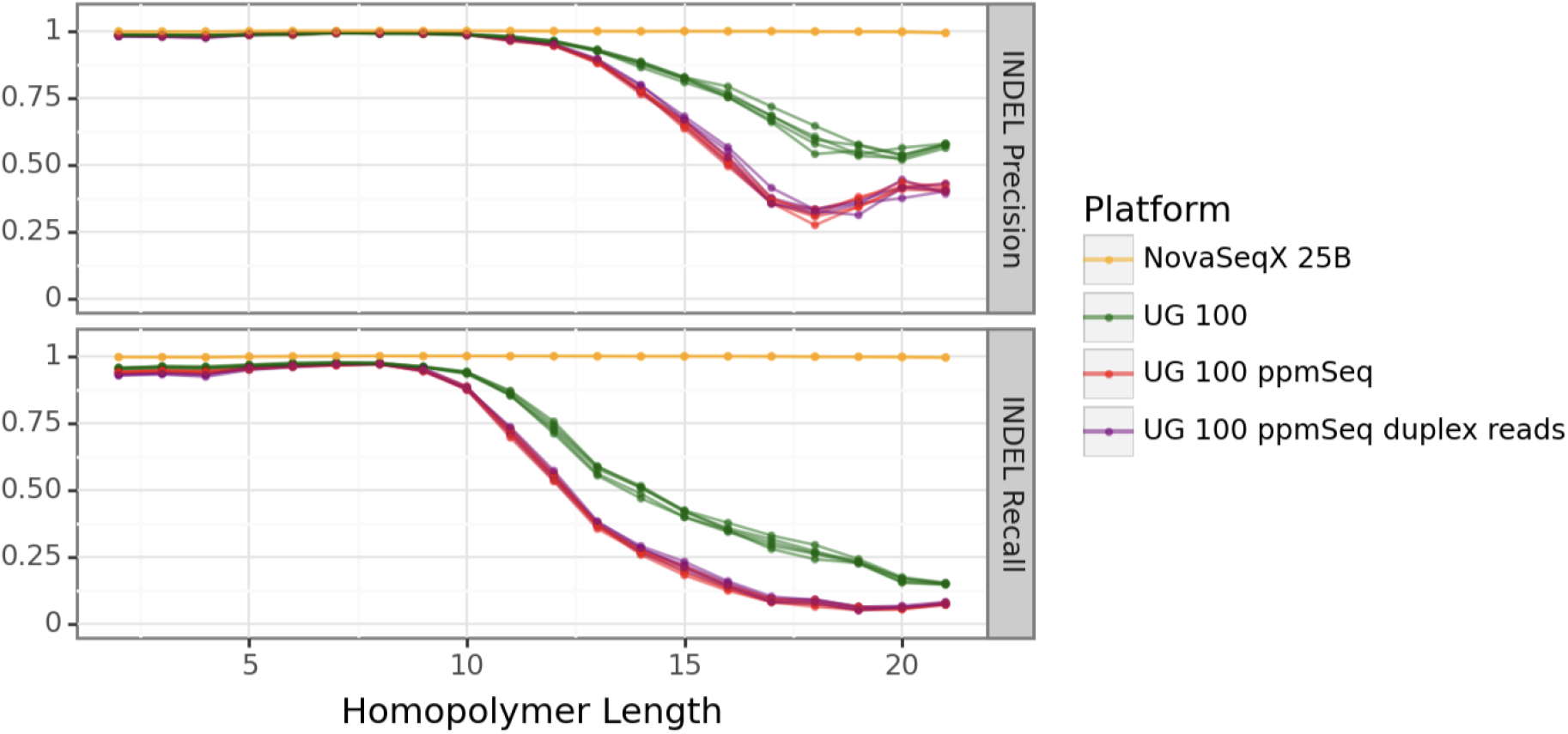
INDEL accuracy vs homopolymer length. Ultima Genomics UG100 variant calling shows a steep drop in precision and recall beyond 10bp homopolymers, reaching precision ranges 0.56-0.58 and recall ranges 0.14-0.15 for homopolymers > 20 bp, while NovaSeqX remains essentially flat across lengths. The trend is more pronounced in UG100 ppmSeq datasets for homopolymers of length 10 and above.

**Figure 4.**
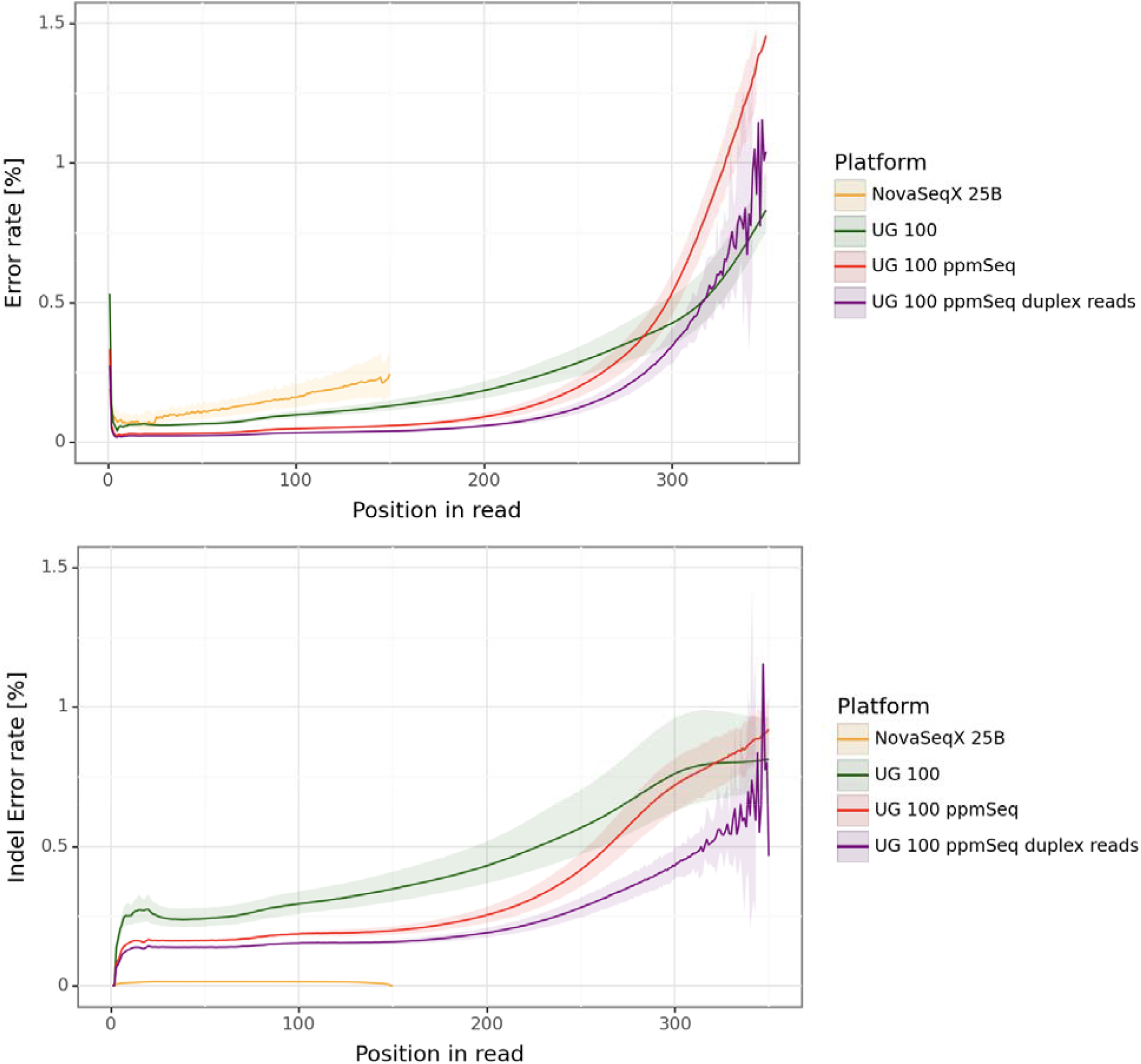
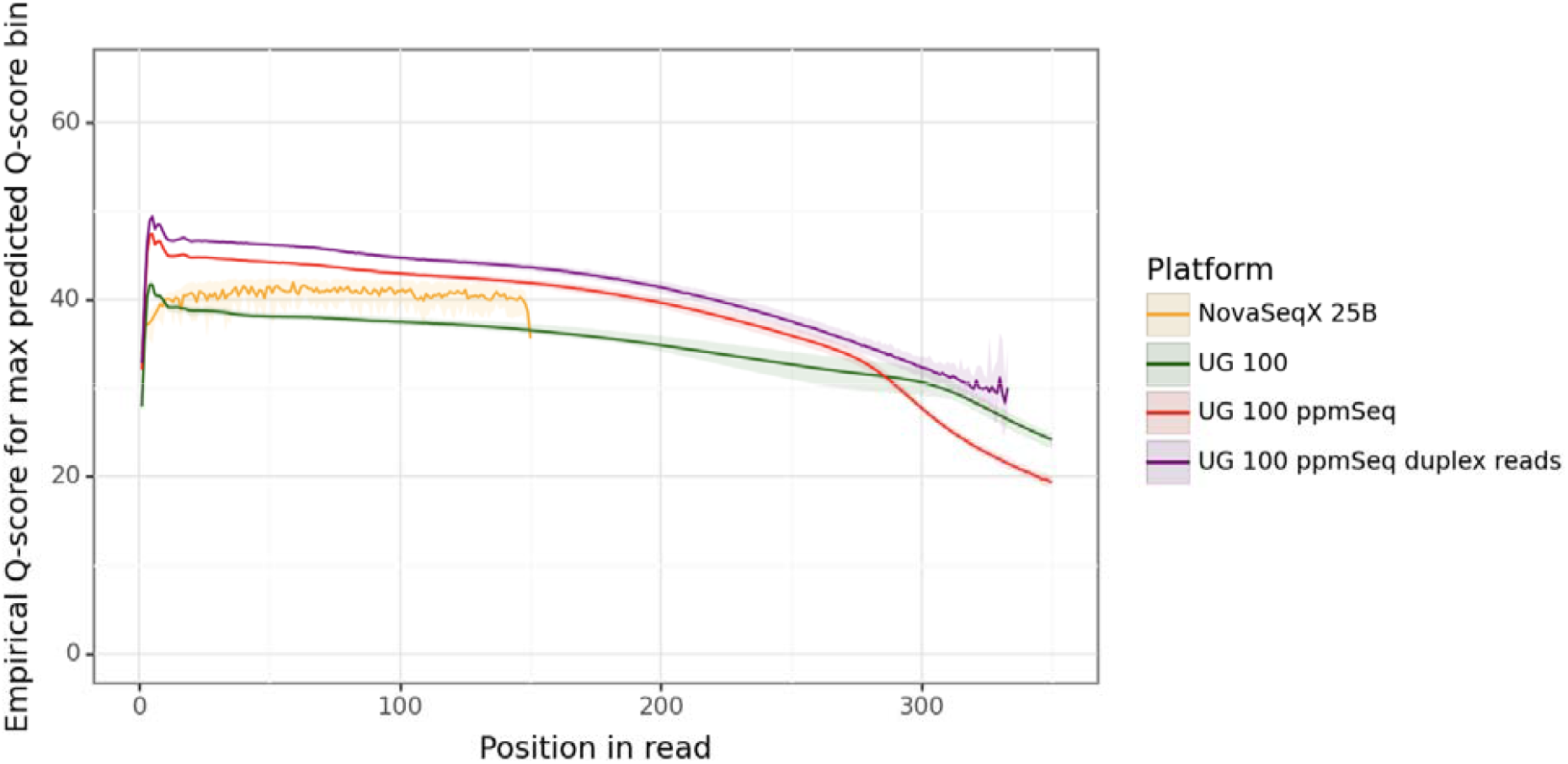
Read-cycle error ramp and indel error is high in UG100. Substitution and indel (combined insertion and deletion) error rates are shown for all platforms and read types up to position 350 in the read. Few UG100 reads exceed 350bp in length. In UG100 reads, substitution error rates stay low early in the read and then climb sharply after position 200 in the read; duplex ppmSeq reduces the levels but the upward slope remains. Indel error rates in UG100 reads are high throughout the whole length of the read. Additionally, substitution errors for all bases that fall in the highest predicted Q-score bin are shown on a phred scale as empirical Q-scores. Highest predicted Q-score bins are Q-40 for NovaSeqX (reported for an average of 94.2% bases) and Q-35 for UG100 (reported for an average of 84.33% in standard chemistry runs and 87.75% in ppmSeq runs).

**Figure 5.**
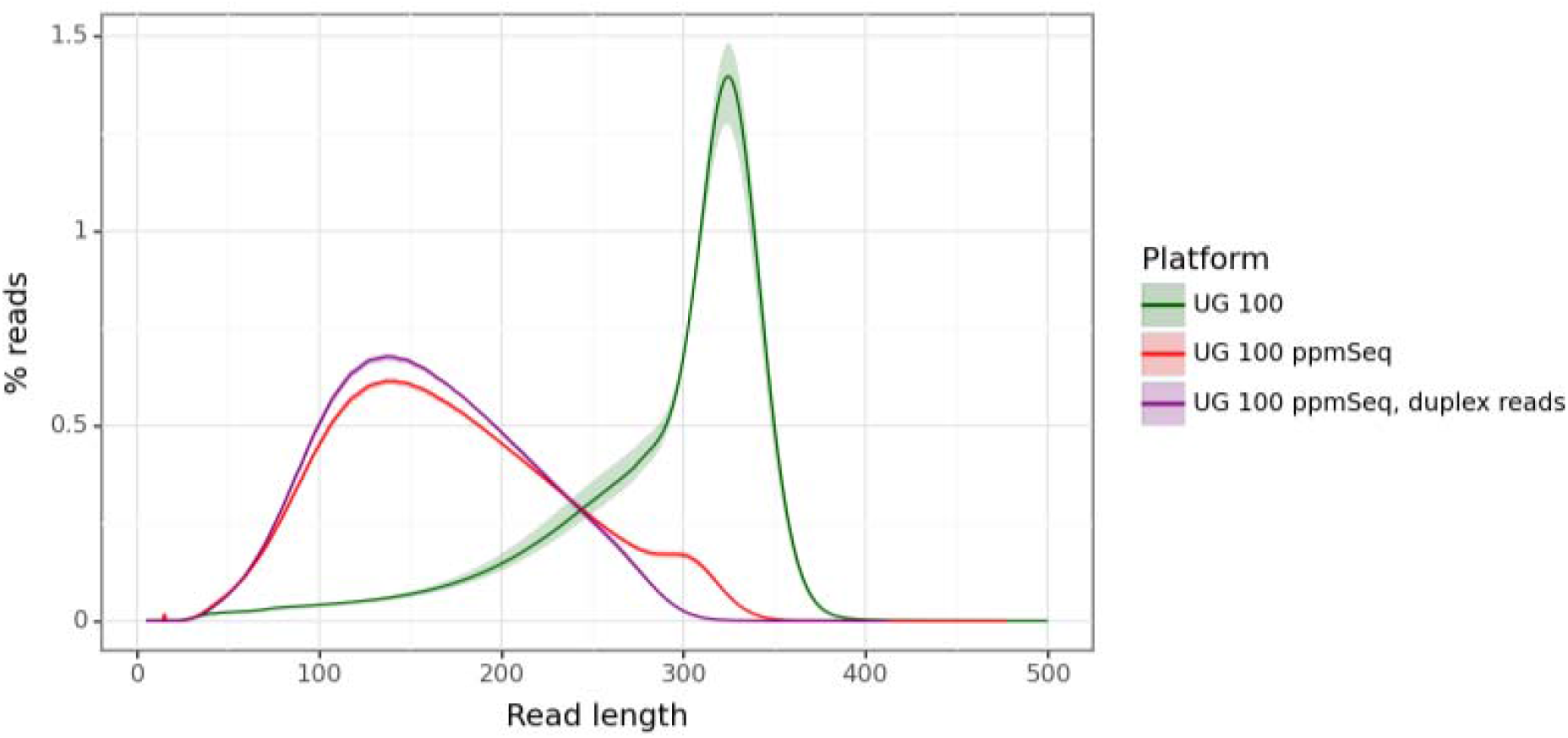
Read□length distributions for UG100 datasets. Read length distributions are shown for all HG002 datasets on UG100 platforms. Solid line indicates median value across replicates, shaded area indicates minimum to maximum range. UG100 median read length is 311bp (median value across replicates), placing many bases in this high-error tail. UG100 read length in ppmSeq runs is shorter, with a median of 165bp in all reads and 157bp in duplex reads only.

**Figure 6.**
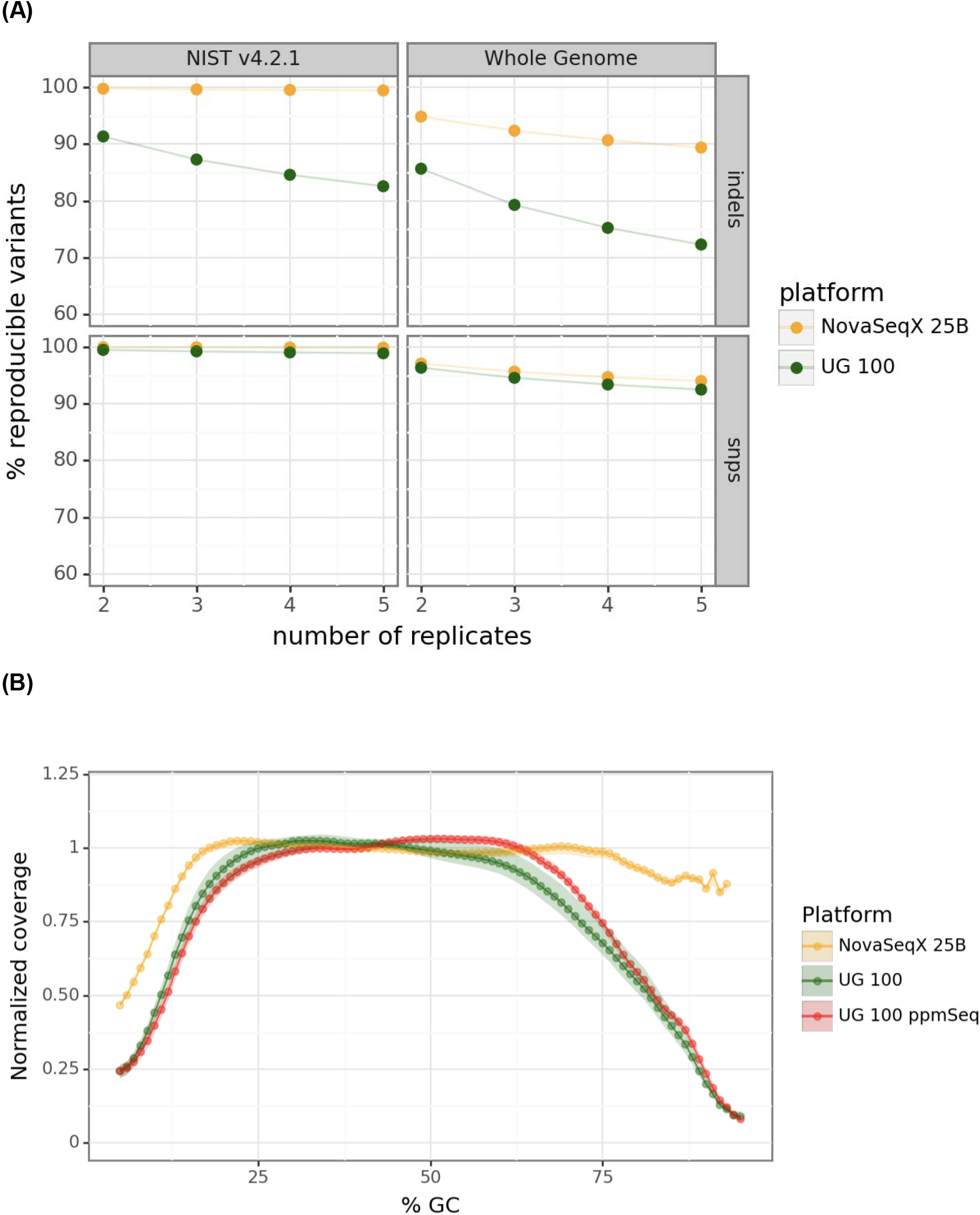
Reproducibility and GC coverage bias. (A)Across HG002 replicates generated from 2 to 5 sequencing runs, NovaSeqX demonstrates higher variant calling reproducibility for SNPs and Indels than UG100. The gap is most pronounced for Indels, both when measured in the whole genome or restricting to NIST v4.2.1 high confidence regions. (B)Normalized coverage by GC content declines sharply in UG100 data beyond 60% GC levels. NovaSeqX 25B normalized coverage profile remains flatter across a range of high GC content.

## Supplementary Note 1

Read-length distribution

Tool: samtools v1.15.1

Command:

samtools stats -@ 8 --cov-threshold 0 --coverage 0,5000,1 --insert-size 20000 --GC-depth 100 - d -p ${cram} > ${sample}.stats

## Supplementary Note 2

Empirical Base-Calling Error Estimation

- We used reads with MAPQ > 30 within NIST v4.2.1 high-confidence regions.
- We masked true variants from the truth VCF to avoid counting real polymorphisms.
- We traversed read CIGAR and compared each read base to reference base; counted total number of bases, mismatches, insertions and deletions.
- Stratified the reported counts by predicted Q-score, read pair (R1/R2), and position in the read.
- Substitution error rates by position in the read were calculated in each replicate by dividing the number of mismatch bases by the number of total bases for each read (R1/R2) and position in the read.
- Indel error rates by position in the read were calculated in each replicate by dividing the number of insertions plus the number of deleted bases by the number of total bases for each read (R1/R2) and position in the read.
- Substitution error rates for the highest Q-score bin were calculated as described above, restricting to bases with a predicted Q-score of Q-40 in NovaSeqX data and Q-35 in UG100 data. They were subsequently reported as empirical Q-scores on a phred scale by applying the following conversion: Q-score = -10 * log10(error_rate)

## Supplementary Note 3

Variant calling benchmark by homopolymer length

- We generated BED file annotations by scanning the fasta file for the human genome reference GRCh38 for consecutive stretches of the same nucleotides of length ≥ 2 bp. We restricted the annotations to autosomes.
- We reported annotations for homopolymers of different lengths (2bp, 3bp, 4bp, …, and > =20bp) in a separate bed file.
- We generate a stratification.tsv file reporting in each row the file name (or any other appropriate label) and the file path (tab-separated) for each homopolymer bed file generated in the step above.
- We benchmarked variant calling by executing the following hap.py script (hap.py v0.3.15):

~~~
  hap.py ${truth_vcf} ${vcf_file_path} \
  -r ${ref_fasta} \
  -f ${truth_confidence_regions_bed} \
  -o ${out_dir}/${out_prefix} \
  --threads 20 \
  --write-counts \
  --engine vcfeval \
  --stratification stratification.tsv
~~~

## Supplementary Note 4

Downsampling and Variant Calling

NovaSeqX 25B and 10B v1.3 datasets

- We processed data from FASTQ by running the DRAGEN v4.4 Germline pipeline ^5^ with “Map/Align + Small Variant Caller” using a GRCh38 graph reference.
- To achieve the target 35x raw downsampling coverage, we:
  - Calculated a downsampling fraction for each replicate as follows: Downsampling fraction = (genome_size × 35) / total_input_bases
  - Ran the DRAGEN pipeline using the following parameters: --enable-fractional-down-sampler=true and --down-sampler-normal-subsample=[downsampling_fraction].
- We additionally ran the DRAGEN pipeline using the --enable-personalization true option UG100 datasets
- We calculated mean coverage in the original aligned CRAMs by:
  - Generating annotations in bed format for GRCh38 autosomes, excluding GRCh38 gaps (GRCh38_autosomes_gaps_removed.bed)
  - Calculated CRAM statistics by running samtools stats restricted to the annotations above and excluding unmapped, secondary and supplementary alignments: samtools stats -@ 8 --cov-threshold 0 --coverage 0,5000,1 --filtering-flag 2820 --target-regions GRCh38_autosomes_gaps_removed.bed ${cram} | grep ^COV | cut -f 2-> coverage_hist
  - We computed mean_raw_coverage by summing the total number of bases (multiplying and summing the second and third columns of the output file) and dividing by the total size of the genome (third column of the output file)
- We calculated a downsampling target fraction for each replicate as 18 / mean_raw_coverage
- We selected 4 libraries of HG002 for each UG100 sequenced run and downsampled each replicate CRAM to 18× by running the following command: ~~~
samtools view -s ${downsampling_fraction_18x} -C -o ${cram_18x} ${cram}
~~~
- For two pairs of libraries on each run, we merged the downsampled CRAMs to generate a 36x merged CRAM. The same libraries were merged together on each run. We achieved this by running the following command: ~~~
samtools merge ${cram1_18x} ${cram2_18x} -o ${merged_cram_35x} -O CRAM
~~~
- Variant calling was performed on the merged CRAM file via the “Ultima Genomics DeepVariant for up to 40x AWS Ready2Run” workflow ID 1617262, version 1.0.

## Supplementary Note 5

Variant Calling Benchmarking stratified by UG100 HCR regions

- We created a UG Low Confidence Regions annotation bed file by taking the complement of the public UG HCR v3.1 bed file. We achieved this by running the following commands: ~~~
cat {ug_hcr_bed} | bedtools sort -i stdin | bedtools merge -i stdin >
ug_hcr_merged_sorted.bed
bedtools complement -i ug_hcr_merged_sorted.bed -g ${ref_fasta}.fai >
ug_lcr.bed
~~~
- We generated a hcr_stratification.tsv file by reporting in each row: 1) an appropriate label and the path to the UG HCR v3.1 bed file (tab-separated) 2) an appropriate label and the path to the UG LCR bed file generated above (tab-separated)
- We benchmarked variant calling by executing the following hap.py script (hap.py v0.3.15):

~~~
hap.py ${truth_vcf} ${vcf_file_path} \
-r ${ref_fasta} \
-f ${truth_confidence_regions_bed} \
-o ${out_dir}/${out_prefix} \
--stratification hcr_stratification.tsv \
--threads 20 \
--write-counts \
--engine vcfeval
~~~

## Supplementary Note 6

Reproducibility Across Replicates

- We selected 5 replicates for each platform to include in this analysis. For UG100, we selected one merged HG002 library (as described in Supplementary Note 5) and the resulting variant calls generated at 35x from 5 sequencing runs. For NovaSeqX, we selected one HG002 library sequenced on 3 sequencing runs on 25B flowcells; additionally, we selected another HG002 library sequenced on 2 sequencing runs on 10B flowcells.
- For each replicate included in the analysis, we extracted PASS SNP and Indel calls from the output 35x VCFs by using the following bcftools command: ~~~
bcftools view -f PASS -v snps ${input_vcf} > $(basename ${input_vcf}
.vcf.gz)_snps.vcf.gz
bcftools view -f PASS -v indels ${input_vcf} > $(basename
${input_vcf} .vcf.gz)_indels.vcf.gz
~~~
- We tabulated inclusion of single variants across multiple replicate variant sets by executing the following command, separately for each variant type (SNP and Indels) and for each platform (NovaSeqX and UG100): ~~~
bedtools multiinter -header -i {vcf_1} {vcf_2} {vcf_3} {vcf_4}
{vcf_5} > ${reproducible_variants_bed_file}
~~~
- We additionally restricted the output bed files to NIST v4.2.1 high confidence regions
- Separately for each variant type (SNP and Indels) and for each platform (NovaSeqX and UG100), we computed the percentage of variant calls that are reproducible in 2, 3, 4, or 5 replicates
  - For example, to calculate reproducibility across 3 replicates, we took all possible combinations of 3 replicates in the set, and for each we calculated the number of variants that are present in 3 replicates and divided it by the number of variants present in > =1 replicate. We then reported an average rate across all possible combinations of 3 replicates in the set.

## Supplementary Note 7

Clinical Impact

- We calculated the percentage of ClinVar pathogenic and likely pathogenic variants in the UG HCR v3.1 regions by:
  - Extracting ClinVar P + LP variants with 2 star and above evidence by downloading the ClinVar variants database and filtering with the following command: ~~~
bcftools view clinvar_{version}.vcf.gz-i ‘
((CLNSIG∼”Likely_pathogenic”) |
(CLNSIG∼”Pathogenic”)) &
((CLNREVSTAT=“criteria_provided” &
CLNREVSTAT=“_multiple_submitters” &
CLNREVSTAT=“_no_conflicts”) |
CLNREVSTAT=“reviewed_by_expert_panel”|
CLNREVSTAT=“practice_guideline”)’ >
clinvar_{version}_P_LP_2_star.vcf
~~~
  - Intersecting the VCF with the UG HCR v3.1 BED file (bedtools intersect)
- We calculated the percentage of overlap with polymorphic short tandem repeats sites by downloading a common catalogue ^8^ and intersecting it with the UG HCR v3.1 BED file (bedtools intersect).

## Supplementary Note 8

Software and Reproducibility

samtools 1.21–1.15.1 ^9^;

bcftools 1.22 ^9^;

bedtools 2.32.0 ^11^;

hap.py 0.3.15 ^10^;

DRAGEN 4.4 ^5^;

AWS Ready2Run DeepVariant v1.0

## Data Availability

Raw sequencing data for all UG100 and NovaSeqX HG002 replicates have been deposited in the NCBI Sequence Read Archive (SRA) under BioProject PRJNA1427896 (SRA submission SUB16023088). The GIAB NIST v4.2.1 truth VCF, high-confidence BED, and HG002 reference materials are available at https://ftp-trace.ncbi.nlm.nih.gov/ReferenceSamples/giab/. Ultima High Confidence Regions v3.1 were obtained from the Ultima Genomics public resources. ClinVar release 2024-08-05 was downloaded from https://ftp.ncbi.nlm.nih.gov/pub/clinvar/.

## Code Availability

All analysis scripts, including hap.py runner scripts, error-by-position pileup code, reproducibility computation, and plotting code are available at https://github.com/oonashigenorisseadams/ug100-vs-nsx (repository will be made public upon data availability in SRA).

## Acknowledgements

OS.R-A. was supported by the Cornell University Graduate School Deans Excellence Fellowship. T.M.N. was supported by a Medical Scientist Training Program grant from the National Institute of General Medical Sciences of the National Institutes of Health under award number: T32GM152349 to the Weill Cornell/Rockefeller/Sloan Kettering Tri-Institutional MD-PhD Program. NGS runs were provided by Illumina.

## Author Contributions

CE, CEM designed the study. OSR-A, CC, and PFP reviewed methodology for analyses. TMN supported analyses. OSR-A and CC wrote manuscript text and figures. All authors contributed to manuscript review.

## Competing Interests

CC, PFP, CE, are employees of Illumina. TMN holds equity in Illumina, Inc. CEM is Co-Founder of Cosmica Biosciences and Biotia.

## Notes

https://github.com/oonashigenorisseadams/ug100-vs-nsx

http://www.ncbi.nlm.nih.gov/bioproject/1427896

## References

1. Zook JM, McDaniel J, Olson ND, et al. An open resource for accurately benchmarking small variant and reference calls. Nat Biotechnol. 2019;37:561–566. doi:10.1038/s41587-019-0074-6

2. Zook JM, Catoe D, McDaniel J, et al. Extensive sequencing of seven human genomes to characterize benchmark reference materials. Sci Data. 2016;3:160025. doi:10.1038/sdata.2016.25

3. Schneider VA, Graves-Lindsay T, Howe K, et al. Evaluation of GRCh38 and de novo haploid genome assemblies demonstrates the enduring quality of the reference assembly. Genome Res. 2017;27:849–864. doi:10.1101/gr.213611.116

4. Poplin R, Chang PC, Alexander D, et al. A universal SNP and small-indel variant caller using deep neural networks. Nat Biotechnol. 2018;36:983–987. doi:10.1038/nbt.4235

5. Behera S, Rossi M, Wang Y, et al. Comprehensive genome analysis and variant detection at scale using DRAGEN. Nat Biotechnol. 2025;43:1177–1191. doi:10.1038/s41587-024-02299-x

6. Almogy G, Pratt M, Bhatt S, et al. Cost-efficient whole genome-sequencing using novel mostly natural sequencing-by-synthesis chemistry and open fluidics platform. bioRxiv. 2022. doi:10.1101/2022.05.29.493900

7. Landrum MJ, Chitipiralla S, Brown GR, et al. ClinVar: improvements to accessing data. Nucleic Acids Res. 2020;48:D845–D852. doi:10.1093/nar/gkz972

8. Dolzhenko E, English A, Dashnow H, et al. Characterization and visualization of tandem repeats at genome scale. Nat Biotechnol. 2024;42:1606–1614. doi:10.1038/s41587-023-02057-3

9. Danecek P, Bonfield JK, Liddle J, et al. Twelve years of SAMtools and BCFtools. GigaScience. 2021;10:giab008. doi:10.1093/gigascience/giab008

10. Krusche P, Trigg L, Boutros PC, et al. Best practices for benchmarking germline small-variant calls in human genomes. Nat Biotechnol. 2019;37:555–560. doi:10.1038/s41587-019-0054-x

11. Quinlan AR, Hall IM. BEDTools: a flexible suite of utilities for comparing genomic features. Bioinformatics. 2010;26:841–842. doi:10.1093/bioinformatics/btq033

12. Cleary JG, Braithwaite R, Gaastra K, et al. Comparing variant call files for performance benchmarking of next-generation sequencing variant calling pipelines. bioRxiv. 2015. doi:10.1101/023754

13. Wagner J, Olson ND, Harris L, et al. Benchmarking challenging small variants with linked and long reads. Cell Genomics. 2022;2:100128. doi:10.1016/j.xgen.2022.100128

